# Anatomical Modeling and Optimization of Speckle Contrast Optical Tomography

**DOI:** 10.1101/2023.09.06.556565

**Authors:** Chen-Hao P. Lin, Inema Orukari, Lisa Kobayashi Frisk, Manish Verma, Sumana Chetia, Faruk Beslija, Adam T. Eggebrecht, Turgut Durduran, Joseph P. Culver, Jason W. Trobaugh

## Abstract

Traditional methods for mapping cerebral blood flow (CBF), such as positron emission tomography and magnetic resonance imaging, offer only isolated snapshots of CBF due to scanner logistics. Speckle contrast optical tomography (SCOT) is a promising optical technique for mapping CBF. However, while SCOT has been established in mice, the method has not yet been demonstrated in humans - partly due to a lack of anatomical reconstruction methods and uncertainty over the optimal design parameters. Herein we develop SCOT reconstruction methods that leverage MRI-based anatomical head models and finite-element modeling of the SCOT forward problem (NIRFASTer). We then simulate SCOT for CBF perturbations to evaluate sensitivity of imaging performance to exposure time and SD-distances. We find image resolution comparable to intensity-based diffuse optical tomography at superficial cortical tissue depth (∼1.5 cm). Localization errors can be reduced by including longer SD-measurements. With longer exposure times speckle contrast decreases, however, noise decreases faster, resulting in a net increase in SNR. Specifically, extending exposure time from 10μs to 10ms increased SCOT SNR by 1000X. Overall, our modeling methods provide anatomically-based image reconstructions that can be used to evaluate a broad range of tissue conditions, measurement parameters, and noise sources and inform SCOT system design.

## 1. Introduction

Cerebral blood flow (CBF) is a crucial biomarker for many brain diseases such as acute stroke and traumatic brain injury.^1,2^ Thus, the development of real-time mapping of CBF is desired in clinics. However, reliable methods to measure real-time CBF at the bedside are not currently available. The dominant CBF imaging methods, including positron emission tomography (PET)^3^ and arterial spin labeling magnetic resonance imaging (ASL-MRI),^4^ offer only snapshots due to scanner logistics. Both modalities are incapable of continuous monitoring of CBF and therefore miss crucial dynamic events such as those involved in acute stroke and traumatic brain injury and dynamics related to cerebral autoregulation and response to neurointerventions. Furthermore, the high instrument cost (> 1 million USD) and large facility size have hindered their capability for bedside clinical applications. Diffuse correlation spectroscopy (DCS) is an optical alternative for monitoring CBF and has begun to show promise as a surrogate for PET and ASL-MRI.^5^ DCS utilizes the dynamics of a single speckle ^6^ and can be used at the bedside for continuous monitoring. However, DCS suffers due to a low signal-to-noise ratio (SNR) because each detector measures only one speckle, which requires keeping the detection collection area extremely small (∼1-4 microns).^6^ The small-collection DCS detection translates into very few photons compared to other deep tissue optical systems like functional near infrared spectroscopy that measure light intensity. Intensity measures commonly use fibers or detectors with areas measured in the millimeters (1-4 mm), resulting in roughly 10^4^-10^6^ more photons than typical DCS.

Speckle contrast optical tomography (SCOT) is an attractive CMOS-camera-based imaging approach for mapping CBF that can overcome the limitations of current CBF techniques.^7,8^ Like DCS, SCOT provides an ionizing-radiation-free, portable, and relatively inexpensive method for mapping CBF continuously, overcoming limitations of PET and ASL-MRI. It also addresses the SNR limitation of DCS by aggregating measures from many speckles at once ^7^ - up to order one million speckles - and provides depth-resolution capabilities of tomography by leveraging a dense set of overlapping measurements as used in high-density diffuse optical tomography (HD-DOT). Previous study has shown the feasibility of SCOT for brain imaging in small animals by demonstrating similar results to functional MRI (fMRI) ^9^. However, the previous SCOT systems have used free-space designs and, as such, are not feasible for large-field-of-view CBF neuroimaging in humans due to both limited focusing ability across the head surface and signal attenuation due to hair.

Discrete optode (i.e. source, detector) SCOT systems could address the free-space limitations by both conforming to the complex shape of a human head and also by the combing action of the optodes through hair. Existing fiber-based HD-DOT systems^10–14^ have been mapping cerebral hemodynamics for more than a decade with their fiber tips forming optodes for sources and detectors. Indeed, HD-DOT imaging has been demonstrated extensively in functional neuroimaging paradigms including visual retinotopy,^15–17^ functional connectivity,^18^ language mapping,^11,19^ and decoding of neural activity.^20^ More recently, fiberless HD-DOT systems have retained the same optode design but with active electro-optics.^17,21–26^ These HD-DOT designs can be leveraged for designing SCOT systems. Recent SCOS studies show that cost-efficient multi-mode fibers (MMFs) are suitable for SCOS/SCOT,^27–29^ wherein speckle statistics are preserved through the MMFs and flow information is relayed, e.g., for pulsatile blood flow.^29^ HD-DOT imaging utilizes a linear forward model^10,11,15,30^ relating changes in oxy-hemoglobin (HbO) and deoxy-hemoglobin (HbR) concentrations to differential diffuse optical measurements, with images of changes in HbO and HbR estimated by regularized inversion of the measurements. Computational packages (e.g. NIRFASTer^31^ and NeuroDOT^11^) compute forward and inverse solutions and include anatomical head models, allowing characterization of optical properties for multiple tissue types. NeuroDOT also includes extensive capabilities for data pre-processing and visualization of reconstructed images that can be used for SCOT development.

In this paper, we developed methods for fiber-based SCOT image reconstruction using an anatomical head model. The computing and simulation pipeline we developed uses the NIRFASTer and NeuroDOT packages and includes an anatomical head model with tissue regions for the scalp, skull, CSF, and cortical tissues. Each region is characterized by its optical absorption and scattering characteristics and its baseline blood flow. The anatomical SCOT methods were used to evaluate imaging performance and to optimize system parameters to maximize SNR. Realistic imaging conditions were modeled by incorporating realistic head structures, prototype imaging arrays, and system-specific measurement-SNR. To investigate SCOT over a range of realistic system parameters, we generated SCOT measurements and images in terms of varying exposure time and source-detector measurement sets, and we incorporated models for speckle and instrumental (including shot, read, and dark) noise. To show the importance of an anatomical head model, we evaluated speckle contrast measures as a function of distance under homogeneous and inhomogeneous tissue conditions. To characterize potential SCOT imaging performance, we evaluated imaging resolution, localization error, depth sensitivity, and SNR as a function of exposure time and source-detector distance. These methods and results provide an analytical pipeline for SCOT, and will guide efficient quantitative design of discrete source/detector SCOT systems.

## 2. Materials and Methods

In this study, we extended current SCOT models to allow realistic head geometry, with multiple tissue types, and source and detector positions that mimic those that would be needed in an anatomically realistic high-density SCOT array. To enable the reconstruction of CBF images, an imaging sensitivity matrix was calculated. We implemented this anatomical modeling in a computational pipeline that simulates SCOT measurements and reconstructions.

### 2.1 Modeling Fiber-Based SCOT - Theory

Speckle is the interference pattern produced when using a coherent laser source, which can be theoretically related to the normalized electric field autocorrelation.^32^ The transport of speckles through tissue can be modeled via the correlation diffusion equation.^33,34^ Speckle contrast can be computed directly from images of speckles.^35^ From images or measurements, speckle contrast squared (κ^2^) is defined as the ratio of the variance of intensity (σ^2^) to the mean intensity squared (⟨*I*⟩^2^) ^7^:

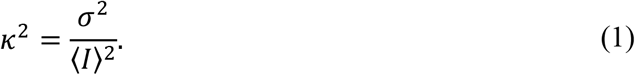

Speckle contrast can also be represented in terms of the normalized electric field autocorrelation, *g*_1_(*r, τ*), in the integral ^7,36,37^:

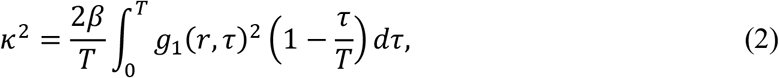

where *τ* is the correlation time, *T* is the camera exposure time, and β is a system constant that depends on the properties of the optical system. The constant *β* is also related to the ratio of the speckle size to the pixel size, e.g., *β* ≤ 0.5 for unpolarized light.^38^

The normalized electric field autocorrelation *g*_1_(*r, τ*) is defined as *G*_1_(*r, τ*)/*G*_1_ (*r*, 0) where *G*_1_(*r, τ*) is the electric field autocorrelation, which depends on optical tissue properties via the well-known correlation diffusion equation ^34,39^:

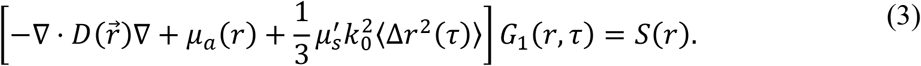

Here, 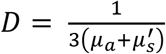 is the photon diffusion coefficient, *μ*_*a*_ is the absorption coefficient, 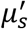 is the reduced scattering coefficient, 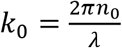 is the modulus of the propagation vector of light, *n*_0_ is the free medium refractive index, *λ* is the wavelength of the laser, Δ*r*^2^(τ) is the mean square displacement of moving particles in the medium, and *S*(*r*) is the photon source.

Following the solution derived by Varma et al.,^8^ equation (3) can be solved to represent a differential speckle contrast measurement, Δ*κ*^2^, resulting from a perturbation 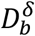 in flow *D*_*b*_ (*mm*^2^/*s*). A general solution can be obtained in terms of baseline Green’s function 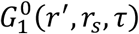, a solution to (3) for an impulsive source at a location *r*_*s*_, *S*(*r*) = δ(*r* − *r*_*s*_), with the measurement at the location *r*^*′*^. Using reciprocity, the detected field can be represented as Green’s function 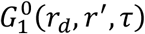 for detector location *r*_*d*_. The general solution includes a factor due to the source and measured at the detector location, given by the ratio of autocorrelation 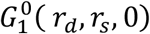 and its normalized version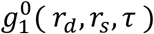. The solution assumes the Born approximation and ignores an external velocity term in the case of in vivo imaging. Here, we have specified positions *r*_*s*_, *r*_*d*_ for the source and detector, respectively, as needed for the anatomical head model, giving differential measurement 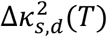 as a function of exposure time *T*:

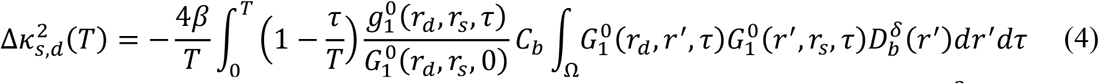

where the coefficient *C*_*b*_ combines multiple terms for simplicity as 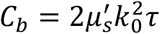. Here we assume < Δ*r*^2^(τ) >≈ 6*D*_*b*_τ as in previous work.^8^ Note that *D*_*b*_ relates to the commonly used blood-flow index (BFI) as BFI = α*D*_*b*_τ, with α, the fraction of dynamic photon scattering events.^37,39^

We can represent eq. (4) as a linear model *y* = *Ax* for measurements 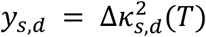, blood flow change 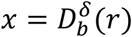 and a sensitivity matrix *A* that maps the change in flow to the measurement:

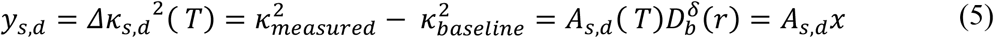

where 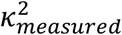 and 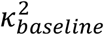 are the measured (perturbation) and baseline values of *κ*^2^, respectively, and explicit dependence on *T* and *r* are not shown in *y*_*s,d*_ = *A*_*s,d*_*x*. We represent the vector of all source-detector measurements as *y*, and the associated sensitivity matrix *A*, resulting in linear model *y* = *Ax*. This model requires a discretized version of the volume integral in (4). The (*i, j*) element of *A* links the i^th^ measurement and the j^th^ voxel as

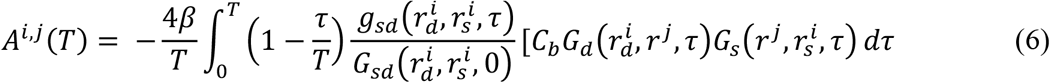

where 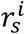 and 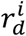 are the locations of the source and detector of the *i*^*th*^ measurement, *r*^*j*^ is the location of the *j*^*th*^ voxel; Green’s functions 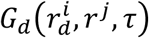 and 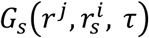 are relabeled as *G*_*d*_ and *G*_*s*_ as the detector and source Green’s functions, respectively; source-detector terms are rewritten as *G*_*sd*_ and *g*_*sd*_; and the dependence of *A* on exposure time *T* is included. For a SCOT array with *N* sources, *M* detectors, and *N*_*x*_, *N*_*y*_, *N*_*z*_ voxels, the matrix *A* is size *NM* measurements by *N*_*x*_*N*_*y*_*N*_*z*_ voxels.

To reconstruct change in flow *x* from the linear forward model *y* = *Ax*, images can be reconstructed as in our previous work in HD-DOT using a generalized Moore-Penrose pseudoinverse with spatially varying regularization ^10,11,30^:

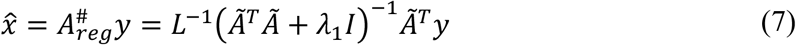

where 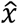 is the reconstructed (estimated) image, 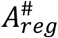 is the regularized pseudoinverse of the sensitivity matrix *A, λ*_1_ is the Tikhonov regularization parameter, and *Ã* = *LA* for *L*, a diagonal matrix given by

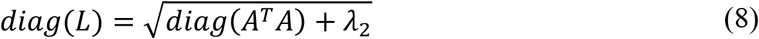

with spatial regularization factor *λ*_2_.

### 2.2 SCOT - Simulation and Image Reconstruction

To further develop and evaluate our SCOT model, we developed a computational pipeline (Fig. 1) that generates simulated measurements and images based on an anatomical head model and a SCOT imaging array. We built upon two existing MATLAB packages: NeuroDOT,^40^ a package for optical brain mapping, and NIRFASTer,^41^ a package for detailed forward modeling. We adapted both packages to compute source and detector Green’s functions and sensitivity (*A*) matrices using an anatomical head model and high-density (HD) source/detector array available in NeuroDOT. We used the linear forward model to simulate measurements for point- and small-volume perturbations in blood flow, and we reconstructed images using existing methods to assess SCOT imaging performance as a function of critical system parameters.

**Figure 1:**
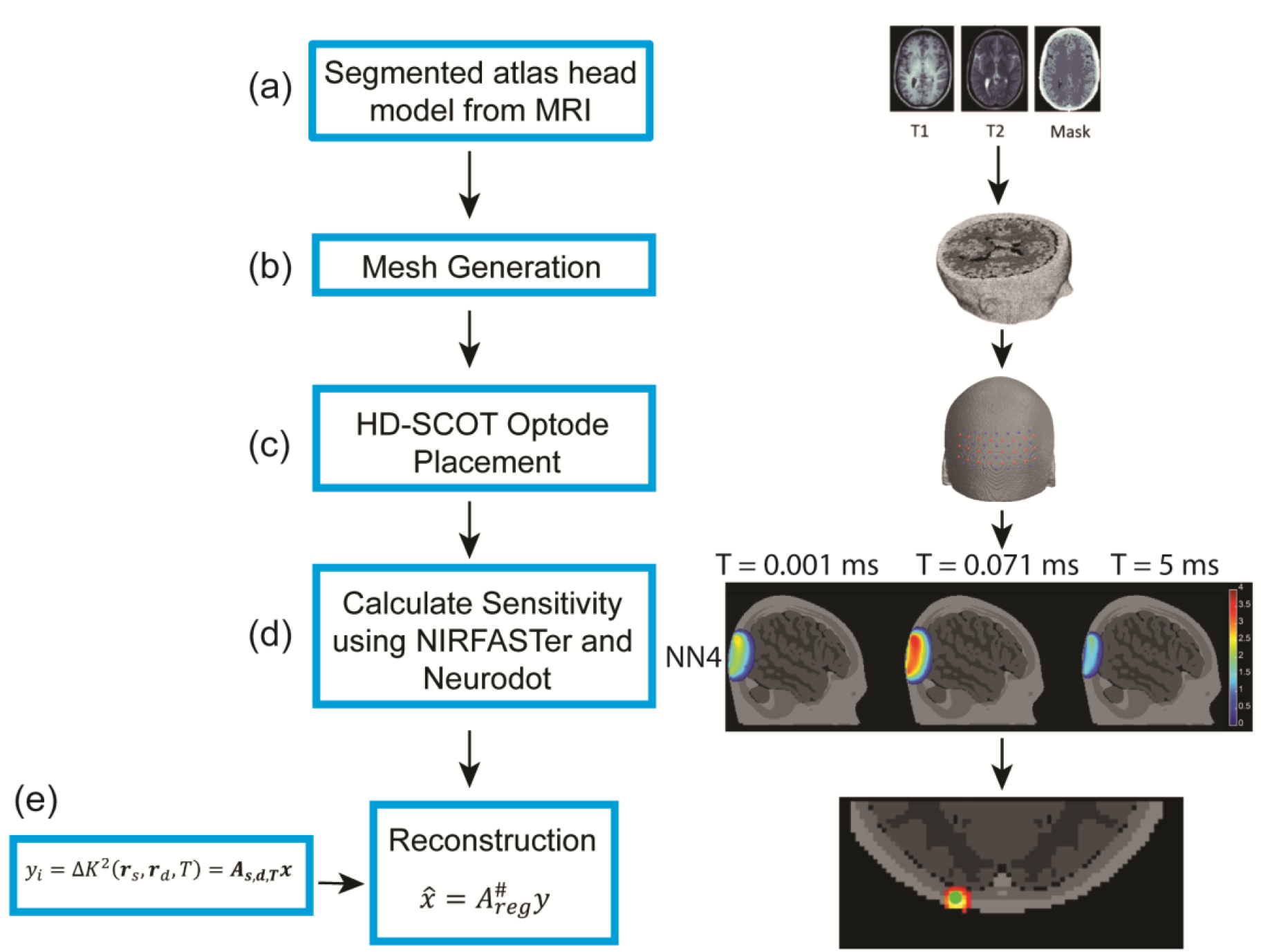
Image reconstruction and simulation methods for SCOT using an anatomical head model. Five tissue types (skin, skull, cerebral spinal fluid, gray matter, and white matter) were segmented from the MNI 152 MRI atlas (a) and used to generate a mesh (b). We positioned 24 source and 28 detector optodes on the head (c) to simulate a high-density (HD) SCOT array. Time-dependent sensitivity matrices for SCOT (d) were simulated using two Matlab packages, NIRFASTer and NeuroDOT. The pipeline is similar to our methods for conventional HD-DOT, with the addition of an exposure time parameter for SCOT imaging. We reconstructed simulated head images (e) using the Moore-Penrose pseudoinverse with spatially varying regularization.

Our forward model produces a sensitivity matrix *A* from an anatomical head model and a set of source and detector positions. Each element of *A* in equation (6) requires correlation-time-dependent solutions for the source and detector Green’s functions. These Green’s functions are computed using the NIRFASTer modeling package based on the optical and flow properties of tissues in the head model. The original NIRFAST^42^ package computes numerical solutions to the diffusion equation for finite-element models (FEM) of an optical medium, allowing more complex geometry than analytic models. The NIRFASTer package extends the FEM capability of NIRFAST with parallelized computation for accelerated processing.^31^ It also provides time-(τ)-dependent solutions to the correlation diffusion equation in (3), e.g., as needed for DCS solutions ^43^ and the Green’s functions of (4). These time-dependent solutions are computed in the NIRFASTer FEM implementation by replacing *μ*_*a*_ in the standard continuous-wave (time-independent) diffusion solution, with the flow- and time-dependent quantity 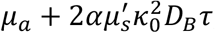, or 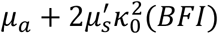. To generate *T*-dependent SCOT sensitivity matrices, we compute Green’s functions using NIRFASTer and then integrate them vs. *τ* as in (6) over the duration of exposure time *T*.

To model SCOT sensitivity matrices for human brains, we adapted a realistic anatomical head model from previous HD-DOT work in our lab,^11^ a tetrahedral mesh based on the MNI152 atlas, with five tissue types (skin, skull, cerebrospinal fluid, gray matter, and white matter), with associated optical properties (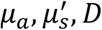, *D*, refractive index *n*, Table 1).^44^ To model background flow in the head model, we considered both homogeneous and inhomogeneous flow conditions. We simulated homogeneous flow ranging from approximately 0.5 to 5 *mm*^2^/*s*, a range of values covering flow rates typical in DCS and SCOT studies.^7,45,46^ We modeled inhomogeneous flow as 1 *mm*^2^/*s* in the superficial regions (scalp, skull, and CSF) and 6 *mm*^2^/*s* in the gray and white matter regions, based on other CBF studies in PET and DCS ^47,48^. Final flow values (Table 1) were validated by comparing simulated decorrelation times (Figure 2) with experimental values for a healthy human subject. We modeled our SCOT array for imaging the visual cortex with 24 sources and 28 detectors, placed on the head at a source-detector spacing of 13 mm using a spring relaxation fitting method (NeuroDOT). Forward model calculations for the array included Green’s functions that vary with *τ* and with source-detector distance *R*_*sd*_, and speckle contrast measurements that vary with *T* and *R*_*sd*_. These results were combined as in (6) to form sensitivity matrices for *T* ranging from 10 *μs* to 10 ms. Sensitivity matrices were visualized for multiple source-detector distances, nominally two-dimensional distances of 13, 29, 39, and 46 mm for first through fourth nearest neighbors, respectively. Forward model speckle contrast results represent baseline conditions. We modeled differential measurements, *y* = Δ*κ*^2^, using the sensitivity matrix in the linear forward model (6) with point-like and volumetric perturbations in blood flow CBF. The resulting data sets included all *NM* source-detector pair measurements as a function of exposure times (10 *μs* to 10 ms).

**Table 1.** Optical and flow properties of tissues in an anatomical head model used for comparing inhomogeneous and homogeneous flow conditions.

| Tissues | Optical Properties |  |  | Inhomogeneous flow | Homogeneous flow 1 | Homogeneous flow 2 |
| --- | --- | --- | --- | --- | --- | --- |
| | | | | $\alpha D_b$<br>[mm<sup>2</sup>/s] | $\alpha D_b$<br>[mm<sup>2</sup>/s] | $\alpha D_b$<br>[mm<sup>2</sup>/s] |
| | $\mu_a$ [mm<sup>-1</sup>] | $\mu_s'$ [mm<sup>-1</sup>] | n | | | |
| Skin | 0.019 | 0.64 | 1.4 | $1 \times 10^{-6}$ | $1 \times 10^{-6}$ | $2 \times 10^{-6}$ |
| Skull | 0.0139 | 0.84 | 1.4 | $1 \times 10^{-6}$ | $1 \times 10^{-6}$ | $2 \times 10^{-6}$ |
| CSF | 0.004 | 0.3 | 1.4 | $1 \times 10^{-6}$ | $1 \times 10^{-6}$ | $2 \times 10^{-6}$ |
| Gray matter | 0.0192 | 0.6726 | 1.4 | $6 \times 10^{-6}$ | $1 \times 10^{-6}$ | $2 \times 10^{-6}$ |
| White matter | 0.0208 | 1.0107 | 1.4 | $6 \times 10^{-6}$ | $1 \times 10^{-6}$ | $2 \times 10^{-6}$ |

**Figure 2:**
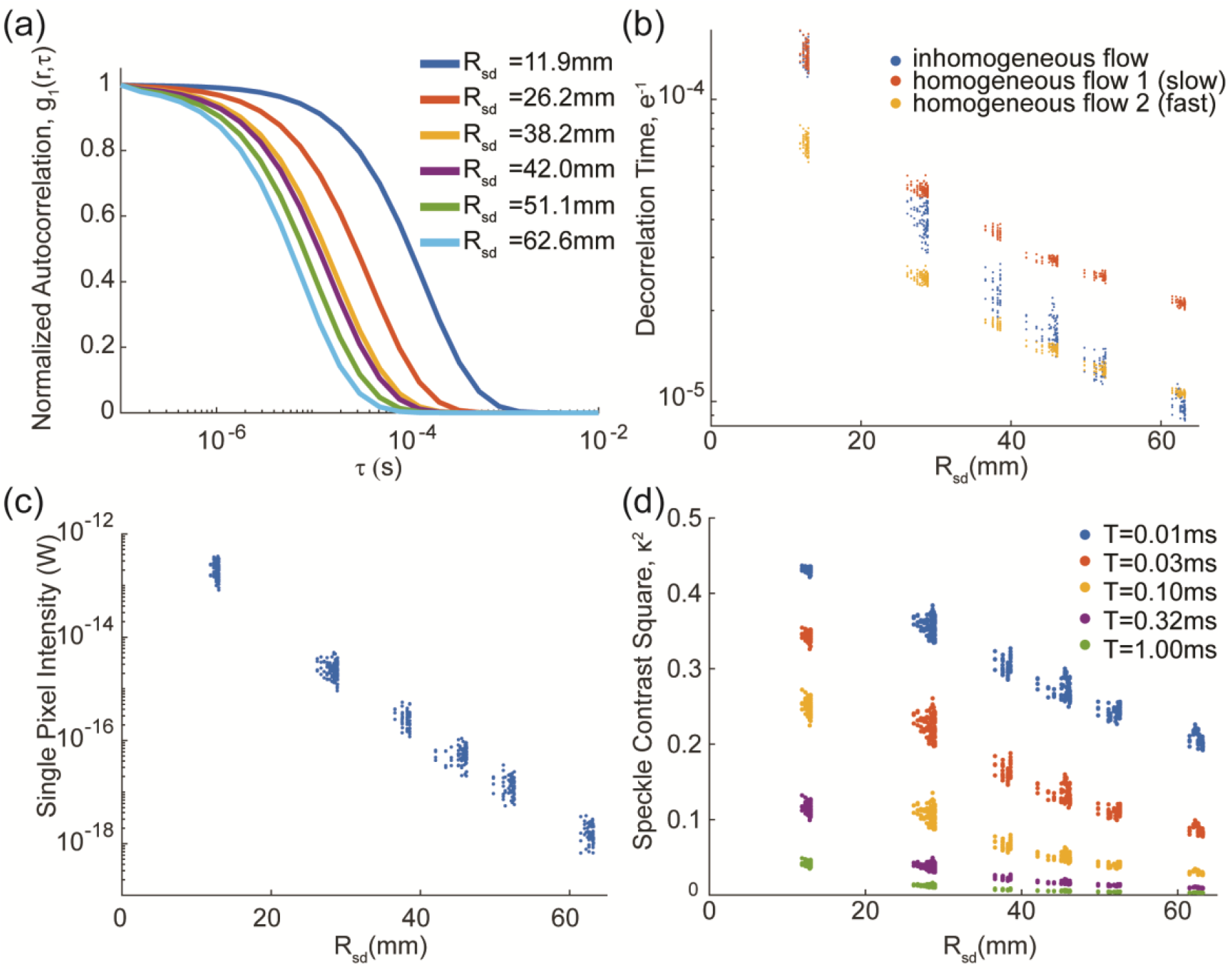
Elements of a simulated SCOT data set using an anatomical head model. Our SCOT sensitivity matrix is built from normalized autocorrelation curves *g*_1_(*r,τ*), (a), with representative curves shown for each nearest-neighbor distance). Decorrelation times (b) were extracted from the autocorrelation curves for all 672 SCOT measurements, and for three flow conditions (two homogeneous, one inhomogeneous). For the inhomogeneous flow condition, simulation data included effective continuous-wave intensity (light falloff curve), (c), scaled to model sCMOS acquisition, and speckle contrast squared, *κ*^2^, versus source-detector distance *R*_*sd*_(d), shown for several exposure times.

To investigate how SCOT system factors affect imaging performance, we simulated images with varied exposure times and source-detector measurement sets. Images were reconstructed from simulated measurements, *y*, as in equation (7), using a generalized Moore-Penrose pseudoinverse and spatially varying regularization. Images were then smoothed with a 3D Gaussian smoothing function (mean of 0 and standard deviation of 3 mm). To evaluate imaging performance, we generated point spread functions (PSF), images of a point-like target, and computed imaging resolution (full-width half-max, FWHM, and full-volume half-max, FVHM), localization error, and effective resolution. The depth of the point-like CBF perturbation was varied, and images were reconstructed from measurements with varying maximum source-detector distance, e.g., all measurements with *R*_*sd*_ ≤ 30 *mm*. Localization error was evaluated as the distance between the centroid of the reconstructed volume and the actual location of the target.^49^ Effective resolution^49^ was characterized as the diameter of the sphere enclosing all voxels with intensity at least half the maximum value.

### 2.3 Noise Models and SNR

To examine how SCOT images are affected by measurement noise or variance, we incorporated multiple noise models and calculated both measurement and imaging signal-to-noise ratio (SNR). Noise was modeled as zero-mean Gaussian data, with variance depending on the source-detector distance, i.e., *y* = *Ax* + *∈*, with 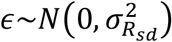. Three different noise models were used.^50,51^ These models were derived from speckle image distributions generated with an adapted copula-based method,^52^ with detailed modeling of the experimental setup (for CMOS camera Orca OrcaFlash4 v3, Hamamatsu Photonics K.K.),^51^ including optical intensity, optical properties of the medium, source-detector distance, exposure time, speckle-pixel ratio, window size, and the number of independent speckles used for computing speckle contrast (parameters, Table 2). Variance 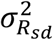 was computed from multiple simulation trials. These values were fitted via linear regression of the log-compressed values as an exponential function of source-detector distance and exposure time for each model. The simplest noise model included variance due only to speckle, i.e., with no instrument noise. The other two models also included system noise, either speckle + shot noise, or speckle + shot + dark + read noise, and both included correction using a theoretical denoising method.^7^

**Table 2.** Parameters for the SCOT simulation.

|  | Parameter | Values |
| --- | --- | --- |
| Noise model parameters | Illumination (photons/s) | ~30 to 330,000 |
| | Exposure time, $T$ | $10^{-5}s$ to $10^{-2}s$ |
|  | Speckle diameter | 3 pixels/speckle |
| | Window size | $100 \times 100$ (pixels) |
| | Wavelength ($\mu m$) | 0.785 |
|  | Number of simulation trials | 100 |
| SCOT System Example design parameters | Optode spacing (mm) | 13 |
|  | Number of sources | 24 |
|  | Number of detectors | 28 |
| | Temporal encoding (time per source) | $\geq T$ |
| | Frame rate (Hz) ($24 \text{ sources} * T$)<sup>-1</sup> | $4 \text{ Hz } (T = 10^{-2}s) \text{ to } 4 \text{ kHz } (10^{-5}s)$ |
| | Detection fiber - core diameter | $200 \mu m$ |
| | Equivalent fiber area | $737 \times 737$ (pixels) |
| | Speckle size ($\mu m$) | 19.5 |
|  | Illumination light levels (mW) per source | 50 mW |
| | Imaging SNR (NN2 at $T = 5 \text{ ms}$) at modeled noise | 0.75 |
|  | Target Imaging SNR, NN2 | 3 |
|  | Increase area/window size | 4x |
|  | Downsampling | 10 Hz to 1 Hz |
| | Imaging SNR - effective | $SNR_{new} = SNR_{old}(\sqrt{4})(\sqrt{10})$ |

To characterize measurement and imaging SNR, measurements and images were simulated for a volumetric perturbation, intended to represent a detectable change in CBF. Measurement SNR was calculated at each source-detector distance as 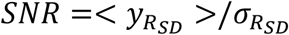, the ratio of the measurement amplitude (signal with maximum amplitude) and the associated noise standard deviation (for that source-detector distance, exposure time, and noise model). For imaging or reconstruction SNR, the signal was determined by summing amplitudes of all voxels greater than 50% of the maximum, and the noise was determined as the mean of the noise standard deviation over the same voxels. Measurements, noise, and SNR were simulated from an exposure time of 10 μs to 10 ms, a significant design parameter that impacts SNR but also limits the possible imaging frame rate (Table 2), e.g., for 24 sources, frame rate ≤ (24 * *T*)^−1^, or 4 *Hz* for *T* = 10*ms*, since sources need to be driven separately to avoid crosstalk.

## 3. Results

The imaging sensitivity matrix in equation (6) is built from Green’s functions describing the diffusion of the electrical field autocorrelation through the head. With an anatomical head model, every source-detector measurement depends on the geometry, flow, and optical properties specific to each tissue type. In contrast, the previous results using an analytic model and a homogeneous medium do not vary for a single source-detector distance. ^50,51,53^

The normalized autocorrelation, *g*_1_(*r*_*d*_, *r*_*s*_, *τ*), varies with each source-detector pair but was visualized (Fig. 2a) for representative measurements at each nearest-neighbor distance (first through sixth nearest neighbors), with the curves denoted as *g*_1_(*r, τ*) with source-detector distance *r* = |*r*_*s*_ − *r*_*d*_|). These results have the form of a sigmoid function, consistent with well-known results in DCS, and the curves vary with increased source-detector distance in agreement with speckle theory.^35^ The decorrelation time (the time for autocorrelation *g*_1_(*r, τ*) to decay to *e*^−1^) relates flow and speckle contrast but also varies with source-detector distance (Fig. 2b). As expected, decorrelation time decreases with longer source-detector distance, since speckles decorrelate more with increased scattering over a longer distance. With the anatomical head model, decorrelation times within each nearest neighbor distance are varied, as expected. In addition to differing tissue paths for each measurement, the actual three-dimensional distance for each pair also varies since the array is fit to an MRI-derived head surface. To demonstrate the potential importance of having different flows in different tissue layers, we simulated measurements with the same (homogeneous) and different (inhomogeneous) flow in the superficial and cortical tissues. The results show a significant difference in decorrelation time between homogeneous and inhomogeneous flow conditions, and the results for inhomogeneous flow with higher flow in deep tissue could not be matched by any homogeneous flow. Furthermore, homogeneous flow at this higher rate has much faster decorrelation (and much lower speckle contrast signal) than when that higher flow is restricted to deeper tissues, suggesting that the normalized electrical field autocorrelation contains important information about the amount of flow at varying depths.

The simulated light intensity (Fig. 2c) decayed exponentially with source-detector distance, as expected from equivalent CW HD-DOT measurements.^11,15,54^ Amplitudes were scaled to reflect single pixel intensity in an sCMOS camera.^54^ Speckle contrast squared (κ^2^, Fig. 2d) was computed as in equation (2) for all measurement pairs and at exposure times from 10 *μs* to 10 *ms*. As expected, speckle contrast decreases with source-detector distance, again due to increased scattering. It also decreases with exposure time due to flow-dependent decorrelation, similar to the inverse relation between speckle contrast and flow rate. Overall, these results (Fig. 2) validate our simulation methods and represent expected variation in SCOT measurements for a realistic head model, under noise-free conditions.

To enable image reconstruction and to understand the depth sensitivity of SCOT measurements to CBF perturbations, we built sensitivity matrices according to equation (6) for multiple exposure times (Fig 3a-c). We visualized the sensitivity matrices at 1 μs, 17 μs, and 4.9 ms over the first (NN1, Fig. 3a), second (NN2, Fig. 3b), and fourth (NN4, Fib. 3c) nearest neighbors (nominal 2D source-detector distances of 13, 29, and 39 mm, respectively). The shape of the SCOT sensitivity is similar to HD-DOT sensitivity, despite an additional scattering term in the correlation diffusion equation, but SCOT sensitivity has a strong and striking dependence on exposure time, with maximum sensitivity around 0.07 ms among these three times. Longer source-detector distances show greater sensitivity to deeper tissues, as in CW HD-DOT, with second and longer nearest neighbors having sensitivity in the superficial cortical region. Thus, one can expect to detect CBF at greater depths with an increased source-detector distance.

**Figure 3:**
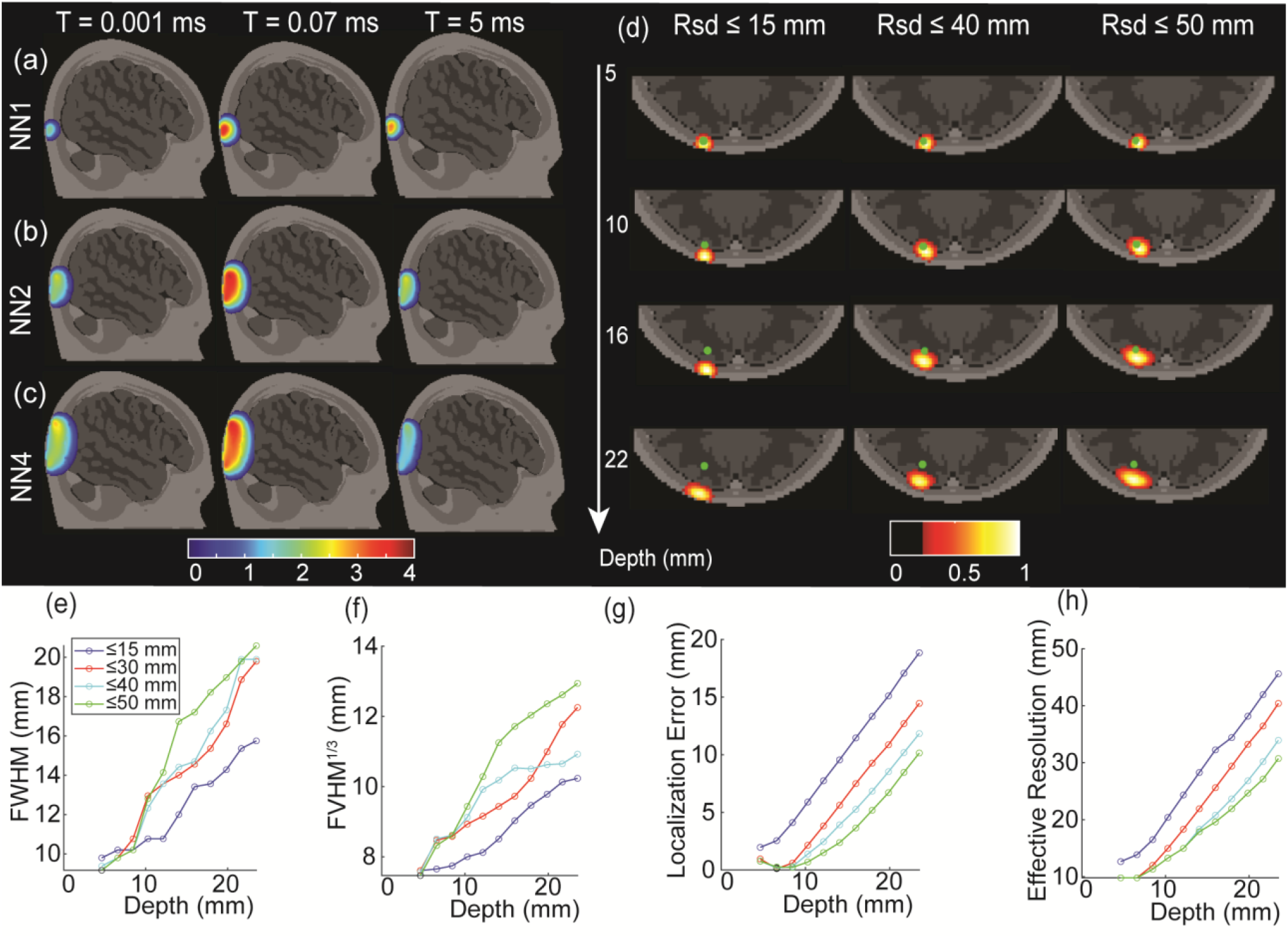
Time-dependent SCOT sensitivity matrices and the reconstruction under various conditions. Short source-detector distance measurements (13 mm, NN1, (a)) show high sensitivity to superficial tissues, while longer source-detector measurements (29 mm, NN2, (b), and 46 mm, NN4, (c)) show higher sensitivity to deeper, cortical tissue. In contrast to HD-DOT, SCOT measurements are highly sensitive to system exposure time, peaking at intermediate times. Intensity is shown on a log scale, with 4 orders of magnitude and scaled to the maximum intensity for each nearest neighbor distance (each row). Reconstructed SCOT point-spread function (PSF, (d)) with varying maximum source-detector distance (*R*_*sd*_*≤*15,40,50 mm) showed the importance of including longer source-detector distances. PSF images were analyzed to assess depth-dependent performance in spatial resolution in full-width half maximum (e), cube root of full-volume half maximum (f), localization error of the reconstructed blood flow (g), and effective resolution (f), again with data sets of varying maximum source-detector distance

To examine SCOT imaging performance, we generated imaging PSFs (Fig. 3d) for simulated measurements from point-like CBF targets at varying depths and using SCOT data sets with maximum source-detector distances of 15, 40, and 50 mm. We analyzed PSFs to estimate image resolution (FWHM, Fig. 3e, and FVHM, Fig. 3f), localization error (Fig. 3g), and effective resolution (Fig. 3h). We observed that including longer source-detector measurements can dramatically improve the localization error. The resolution and effective resolution (Fig. 4b, c, d) were larger (worse) with deeper perturbations. Moreover, the simulation results showed that a SCOT system can achieve a resolution of approximately 13 mm FWHM at 10 mm depth, e.g., deep enough to measure the cortical tissue, similar to results for HD-DOT.^49^ We simulated images for varying exposure times but found negligible variation in these metrics.

**Figure 4:**
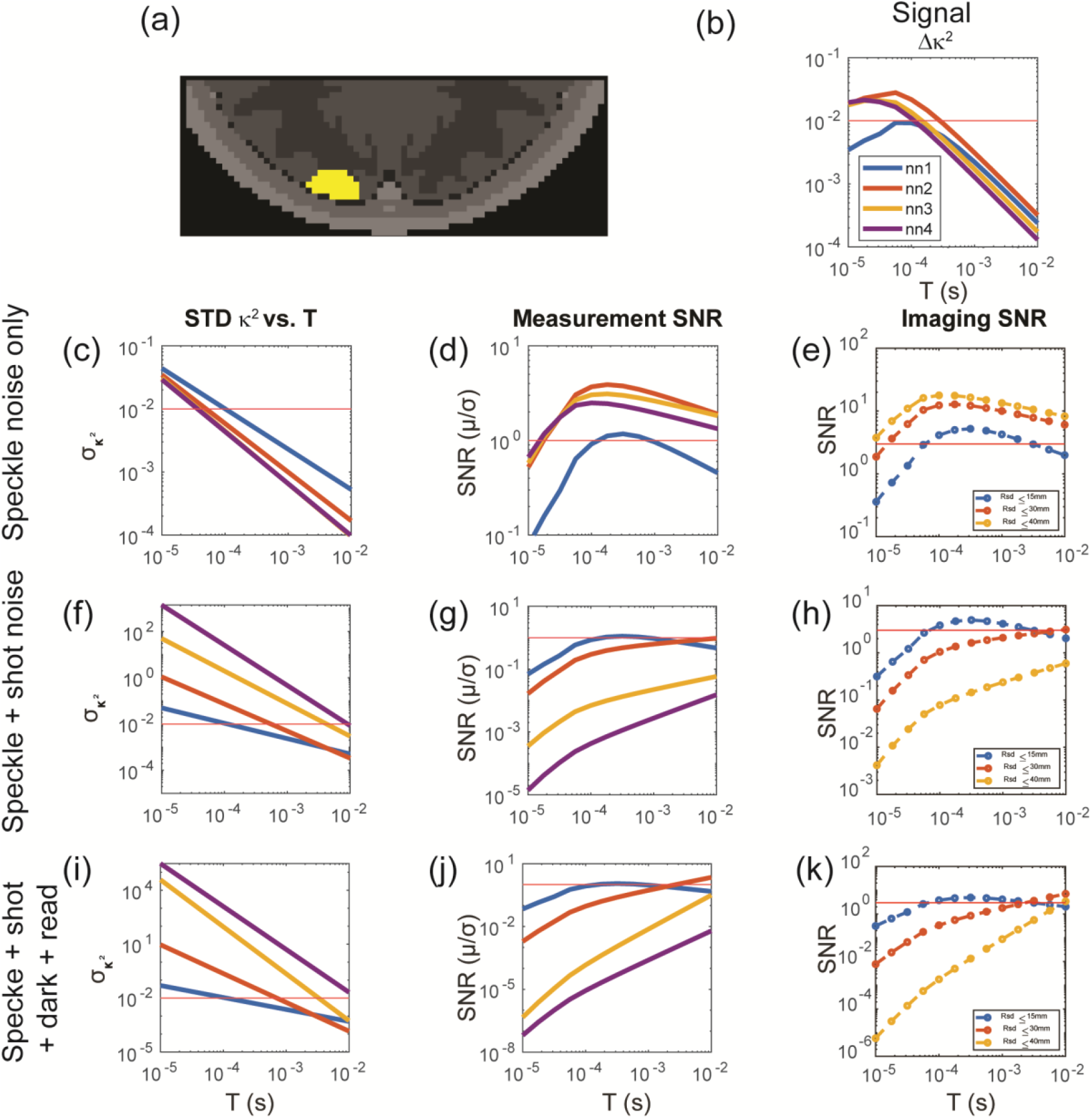
Evaluation of SCOT performance versus noise and exposure time, for data sets with varying maximum source-detector distance (*R*_*sd*_*≤*15,30,40 mm). SCOT measurements and images were simulated for a volumetric CBF perturbation (40% increase in flow) (a), resulting in differential SCOT measurements Δ*κ*^2^ (b). Three increasingly inclusive SCOT noise models (speckle noise only (c,d,e), speckle + shot noise (f,g,h), and speckle + shot + dark + read noise (i,j,k)) were incorporated, allowing calculation of SNR in both measurements and reconstructed images. Noise models characterized baseline variance in κ^2^ as a function of exposure time *T* (c,f,i). Measurement SNR (d,g,j) and imaging SNR (e,h,k) generally increased with *T* and decreased with *R*_*sd*_. For the more inclusive noise models, SNR plateaued around *T =* 5ms and was maximized with *R*_*sd*_*≤*30*mm*. Imaging SNR values assume a 2x increase in κ^2^ window area (relative to noise model assumptions) and a 1 Hz frame rate. The red lines in (b,c,f,j) were plotted at y = 10^−2^, the red lines in (d,g,j) showed the measurement SNR at 1, and the red lines at (e,h,k) represented the target imaging SNR of 3.

To investigate the influence of noise on SCOT measurements and images, we incorporated three different models of noise and characterized measurement and voxel SNR for a volumetric CBF perturbation in the cortical tissue (Fig 4a). We used a copula-based noise model (Kobayashi-Frisk et al.) and characterized noise as variance in the differential speckle contrast measurement versus exposure time and for multiple source-detector distances (Fig. 4c, f, i) for each noise model. We simulated noise-free SCOT measurements (Fig. 4b) and images first, then we added noise to the measurements, and, finally, reconstructed images from those noisy measurements. We computed SNR for both the measurements (Fig. 4d, g, j) and images (Fig. 4e, f, k). In calculating the image SNR, we made two changes to the original parameters in Table 2 to illustrate achieving a target imaging SNR of 3: increase the window size by 4x (still within the detection area of the fiber), and downsample the measurements from 10 Hz to 1 Hz, resulting in 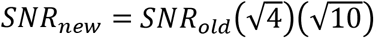, giving a max SNR greater than 3 in these simulations. The first noise model considered speckle noise only (Fig. 4c, d, e), with variance due to the calculation of speckle contrast spatially over a finite window size. We found that, with increasing exposure time, the measurement and imaging SNR both increase to a peak and then decrease, as might be expected from similar results for imaging sensitivity.

In addition to speckle noise, system noise will influence measurements, thus we included two other models to include instrument noise, with speckle + shot noise (Fig. 4f, g, h), and speckle + shot + dark + read noise (Fig. 4i, j, k)), both applied with known noise corrections. In these models, however, measurement and imaging SNR with longer source-detector measurements (NN2 or greater) continued to increase with exposure time up to 10 ms, the maximum we would expect to consider without adversely affecting the overall imaging frame rate.

To visualize the combined effects of exposure time and source-detector distance, we simulated images (Fig. 5) for the volumetric CBF target, in the presence of additive noise (speckle + shot + dark + read noise, Fig. 4i,j,k). Images shown are representative of each condition (individual images vary with each realization of additive noise). We observed that under a short exposure time (0.1 ms), imaging noise appears as artifacts separate from the CBF target. When using longer source-detector distances at this short exposure time, these noisy artifacts are even more apparent, although the location of the reconstructed CBF is closer to the original target. For longer exposure time (10 ms), images still have high noise when formed from only short (NN1) or longer (up to NN3) measurements, but for measurements up to NN2 (*R*_*sd*_ ≤ 30 *mm*) the images have both low localization error and low discernible noise, resulting in a clear view of the CBF target.

**Figure 5:**
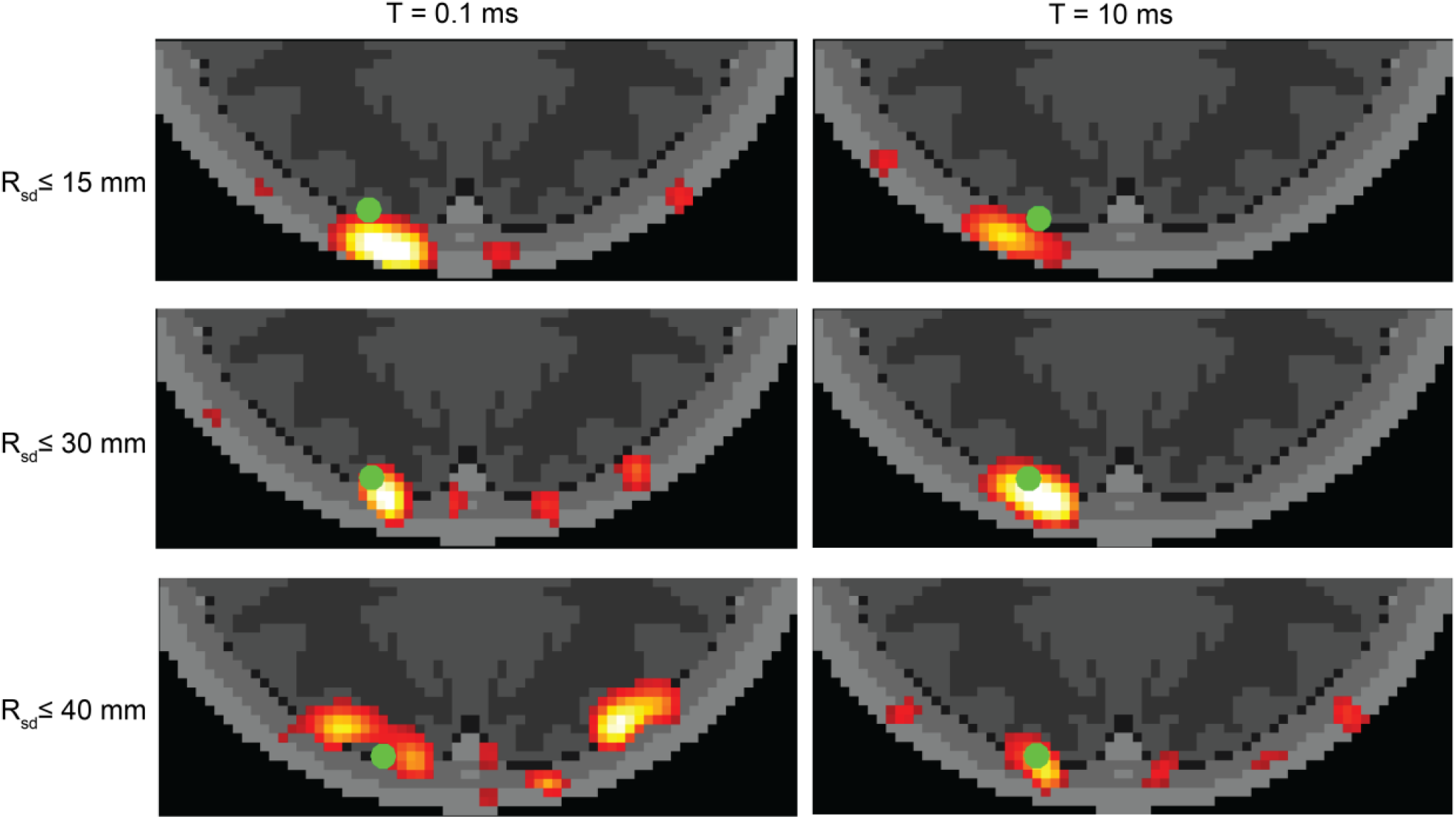
Simulated SCOT images vs. exposure time and source detector distance. Images were reconstructed from data sets with varying maximum source-detector distance (*R*_*sd*_*≤*15,30,40 mm) and with two exposure times (*T =* 0.1, 10 ms) to demonstrate the variance of imaging SNR over a range of possible settings in a SCOT instrument. Images were simulated for the volumetric CBF perturbation (green dot) as in Fig. 4a.

## 4. Discussion

The potential of SCOT for mapping CBF in the human brain will depend on the design and implementation of SCOT imaging systems. Herein we developed an anatomically realistic head model that enables simulation of SCOT measurements. To evaluate imaging of CBF, we extended methods for reconstructing SCOT images and analyzed resolution and localization error, and their dependence on a high-density SCOT measurement set. To understand the impact of system noise on the measurements and images, we incorporated SCOT-specific noise models and evaluated measurement and imaging SNR and the impact of system exposure time. Finally, the SCOT modeling shows that inhomogeneous properties and flow cannot be accurately represented with homogeneous models, with homogeneous models failing at either short or long distances.

Existing models for SCOS and SCOT represent speckle contrast in terms of optical properties and flow within a medium and simulated measurements incorporate relevant noise sources.^8,50^ Forward analytical models for SCOT measurements - with limited analytic, closed-form models ^7,8,10^ such as using a semi-infinite approximation for solving the diffusion equation - are well established. However, the analytic solutions cannot accommodate the complexity of real-world head anatomy. They do not allow realistic surface geometry or complex internal tissue types or varying flow within different tissue compartments. Existing methods for HD-DOT image reconstruction (inverse solutions) have been used for SCOT ^8,10^ but have not been extended to anatomical models. However, recent studies have extended SCOS analysis to include noise sources. Dynamic speckle models incorporating shot, dark and read noise have been developed and demonstrate the variance of speckle contrast statistics with varying intensity, speckle/pixel ratio, and exposure time.^53^ A copula-based simulation model has been developed for modeling speckle dynamics incorporating shot, dark, readout and speckle noise and this model has demonstrated accuracy and precision of speckle contrast calculations, with studies of the variance depending on source-detector distance and exposure time. ^50^,^52^ These recent noise models have significantly improved our understanding of variance in SCOS measurements and could in principle be incorporated into anatomically-based SCOT studies.

Building on previous work demonstrating free-space SCOS/SCOT imaging in simulation and *in vivo* ^7,8,37^, we extended existing theory to develop methods for SCOT modeling and imaging in human brains using a realistic head model and finite element model (FEM) calculations. We modeled measurements for a high-density SCOT imaging array positioned on an anatomical head model with flow and optical properties specified for five distinct tissue types (Fig. 1). FEM-based computational solutions to this inhomogeneous model produced measurements that were sensitive to the geometry, flow, and optical properties of the medium. Simulation of high flow (6 *mm*^2^/*s*) in deeper, cortical tissues and low flow (1 *mm*^2^/*s*) in superficial tissue produced decorrelation times dependent on flow and source-detector distance (Fig. 2), which could not be replicated with homogeneous models as used in previous studies.^7,8,53^ Overall, our forward model for SCOT permits the construction of sensitivity matrices for differential, fiber-based SCOT imaging of CBF, improving on previous free-space designs,^7,37^ and using FEM solutions with greater computational efficiency than alternative photon-based Monte Carlo models.^53^

To evaluate imaging of CBF, we implemented methods for reconstructing SCOT images, based on similar methods for DOT, but with sensitivity matrices strongly dependent on exposure time for SCOT (Fig. 3a-c). We showed that fiber-based SCOT imaging can be expected to achieve a resolution of approximately 1.5 cm for depths at and beyond the superficial cortical gyri (Fig. 3e), comparable to DOT imaging using the same source-detector spacing ^11,49^. We also found that a localization error of approximately 5 mm can be obtained for a CBF perturbation at 10 mm depth, if source-detector measurements up to 30 mm are included, with error reduced further by including longer source-detector distances. Although previous work has demonstrated SCOT imaging in flow phantoms and rodents^7,37^ using free-space instruments and models with analytical functions, our findings establish the first imaging performance expectations based on an anatomical head model for fiber-based SCOT in human brains.

To demonstrate the importance of noise in the overall performance of SCOT imaging, we incorporated system-specific noise models^50^ and analyzed measurement SNR and imaging SNR (Fig. 4). We showed that both depend significantly on exposure time, with measurement SNR increasing ∼1000x from 10 *μs* to 10 *ms* for a measurement with source-detector distance 29 mm, and imaging SNR increasing ∼1000x from 10 *μs* to 10 *ms* when including measurements ≤ 30 *mm*. These results were tailored to a specific noise model, but are consistent with published results^53^, and our methods can easily accommodate other models, e.g., for variations in system design via simulation, or for empirically derived models from actual measurements.

We expect to use our model and simulations to guide the design of future fiber-based SCOT system development, with inclusion of system-specific noise. Our simulations modeled an example system (Table 2) with 24 sources and 28 detectors at 13-mm source-detector spacing, suitable for imaging the visual cortex. With temporal encoding of sources, the imaging frame rate would be limited in a tradeoff with the exposure time, *T*, e.g., with 24 sources, a frame rate of 10 Hz would require 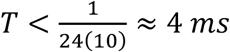. Our simulations showed an imaging SNR of 0.75 for a 40% increase in flow over a cortical region of volume equivalent to a sphere of radius ∼8 *mm*. Imaging SNR ≥ 3 could be achieved by, for example, increasing the speckle window size by a factor of 4 and downsampling from 10 Hz to 1 Hz. In general, measurement SNR depends on the number of speckles measured, thus larger detection fiber is desirable, although at the expense of requiring a greater imaging sensor area. These simulation results suggest a feasible design for fiber-based SCOT imaging of CBF that leverages both previous designs for HD-DOT systems ^11,54^ and free-space SCOT systems ^7^.

Overall, we expect SCOT imaging performance to improve with longer exposure time and longer source-detector distances (Fig. 5). This is not too surprising. While the contrast is higher at shorter exposure times, in the end, the increased exposure times win out, allowing more photons to be collected by the system, and reducing noise (standard deviation). Better localization for CBF changes at deeper depths through the inclusion of longer source-detector measurements is also natural to expect. Our results quantify this relationship. We expect a tradeoff between exposure time and frame rate, and we note that our current results show decreased SNR with measurements > 30 mm. We expect inverse solutions that include the noise variance ^55^ might improve the imaging SNR while allowing the improved localization associated with longer measurements. While our noise models were rigorously developed,^50^ the results are specific to instrument details, and additional work remains to validate these models.

## 5. Conclusion

In conclusion, we have introduced methods for fiber-based SCOT imaging on an anatomical head model. We have simulated imaging performance using a high-density array and with varying system characteristics, including exposure times. Our measurement simulations agreed with previous results in homogeneous media but also showed that results for inhomogeneous media could not be replicated by homogeneous models. We showed that SCOT imaging can be sensitive at the depth of the superficial cortical gyri, with comparable spatial resolution to CW HD-DOT.

We confirmed and quantified that including longer source-detector distances can reduce the localization error but may be limited by system and speckle noise. Overall, the results demonstrate that the exposure time of a SCOT system is a key factor in its performance. A longer exposure time can reduce the signal strength of measurements, but also reduces noise, thus a longer exposure time appears to increase SNR. At exposure times greater than 10 ms, the tradeoff with frame rate can become significant. Overall, our simulations enable anatomically-based SCOT image reconstruction, and can be used to guide the design of fiber-based SCOT system development.

## 6. Acknowledgments

The project was funded by National Institute of Neurological Disorders and Stroke (R01NS090874). We also thank the fellowship and scholarship from ICFO Fundació CELLEX Barcelona, Fundació Mir-Puig, Agencia Estatal de Investigación, (PHOTOMETABO, PID2019-106481RB-C31/10.13039/501100011033, PLEC2022-009290/SAFEICP; MEDLUX; <u>PRE2018-085082</u>), the “Severo Ochoa” Programme for Centres of Excellence in R&D (CEX2019-000910-S), Generalitat de Catalunya (CERCA, AGAUR-2022-SGR-01457, RIS3CAT-001-P-001682 CECH), TV3 La Marato, European commission (Marie Curie – <u>847517, 860185/PHAST; VASCOVID; TINYBRAINS)</u>, ISCIII (FRONTSTAGE; LITEMUSCLE).

## 7. Author Contributions

C.H.P.L., I.O., T.D., J.P.C., and J.W.T. conceived and designed the experiments. C.H.P.L., I.O., L.K.F., and J.W.T. performed the experiments. C.H.P.L., I.O., L.K.F., T.D., J.P.C., and J.W.T. analyzed the data. C.H.P.L., I.O., L.K.F., A.T.E., and J.W.T. wrote analysis software. C.H.P.L., I.O., J.P.C., and J.W.T wrote the manuscript. All authors edited and reviewed the manuscript.

